# Snail host parental investment throughout a *Schistosoma mansoni* infection

**DOI:** 10.1101/2020.11.05.370510

**Authors:** Stephanie O. Gutierrez, Olivia J. Lockyear, Dennis J. Minchella

## Abstract

Parental investment theory describes the ability of organisms to respond to an environmental challenge by increasing the fitness of future offspring. Utilizing life history changes, organisms can maximize fitness by increasing their total reproductive output or by investing more into the success of fewer offspring. In cases where parasitic infections result in castration of their host, increased reproductive effort known as fecundity compensation has been demonstrated in a variety of organisms. This response appears predictive of expected future reproductive losses. Organisms struggling with an environmental pathogen, may attempt to better prepare their offspring for the environment they are experiencing through transgenerational immune priming (TGIP). In immune priming, primary infection lowers the prevalence and intensity of a subsequent infection by the same pathogen. Transgenerational immune priming carries pathogen resistance into further generations without genotypic changes. The focus of this study was to determine whether invertebrate parental investment into offspring parasite resistance varies over the course of an infection. Utilizing the snail host *Biomphalaria glabrata* and its trematode parasite *Schistosoma mansoni*, offspring were reared from specific time intervals in the parent’s infection and subsequently exposed to the same pathogen when each cohort reached the same age- 12 weeks. Differences in infection prevalence and intensity were expected based on when the offspring were born during their parent’s infection. A trade-off was predicted between the number of offspring produced in a cohort and offspring resistance to future infections. Offspring born during the period of fecundity compensation were predicted to exhibit lower resistance due to a dilution of individual investment by parents into a larger offspring pool. While our results did not support TGIP, there were differences in offspring prevalence, as well as an indication that parent health may interact with genetics in offspring resistance. Results suggest that parental condition can influence resistance of *B. glabrata* offspring to *S. mansoni* but that TGIP may not be operating in this system.

---

Resource limitations as well as environmental burdens can impact an organism’s ability to maximize reproduction and result in trade-offs between life history traits. For instance, *Daphnia magna* infected with *Pasteuria ramosa* can trade infection tolerance for an ability to maximize reproductive output (Vale and Little 2012). This life history response is termed fecundity compensation, as exposed hosts have a burst of early reproduction to compensate for the expected future loss of reproductive success (Minchella and Loverde 1981). Fecundity compensation can be seen in cases where parasites ultimately end a host’s ability to reproduce once infected, as hosts must choose between maximizing their defense against the parasite or maximizing their reproduction prior to castration by these parasitic castrators.

*Schistosoma mansoni* is a parasitic castrator in its intermediate snail host and a disease agent in humans. Schistosomiasis, a parasitic disease caused by *Schistosoma spp.,* afflicts more than 250 million people worldwide and remains endemic in parts of Africa, Asia, and South America (Sokolow et al. 2016). Humans are infected by free-swimming cercariae released by infected snails in water sources. Subsequently, the parasite is recirculated into snail populations by infected human waste matter reaching freshwater sources, where miracidia can hatch and infect *Biomphalaria* spp. snails. Understanding snail host life history responses to parasites can provide a better understanding of host-parasite relationships.

Organisms struggling with an environmental pathogen, may attempt to better prepare their offspring for the environment they are experiencing. Immune priming occurs when a primary infection lowers the prevalence and intensity of a subsequent infection by the same pathogen. The occurrence of immune priming in invertebrates challenges a long-standing assumption that invertebrate immunity is non-adaptive and without memory. In the *B. glabrata* and *S. mansoni* system, snails vaccinated with miracidium protein extracts prior to parasite exposure exhibited a lower prevalence of infection and lower infection intensity (Portela et al. 2013). Furthermore, the innate immune memory in *B. glabrata* has been determined to be a humoral immune defense response rather than one based in cellular mechanisms (Pinaud et al. 2016). This within generation immune priming is widespread in invertebrates, and there is growing evidence that invertebrate immune priming can be transgenerational (Roth et al. 2018).

Transgenerational effects characterize the altered offspring phenotypes generated by parents exposed to an environmental stimulus. Transgenerational immune priming (TGIP) thereby carries changes in immunological function across generations without genotypic changes. For example, in *Daphnia magna,* maternal and offspring environments were both found to be major determinants of disease susceptibility. Parental investment into offspring immunity differed depending on the mother’s developmental environment, as offspring from stressed mothers were more resistant than offspring from unstressed mothers (Mitchell and Read 2005). While TGIP has been well documented in vertebrates with the maternal transfer of immune factors which reduce offspring susceptibility to disease, research of TGIP in invertebrates has been more limited (Littlefair, Laughton, and Knell 2017).

The focus of this study is to determine whether a snail’s parental investment into offspring immunity changes over the course of a *S. mansoni* infection. In the infected generation of parent snails, we expect to observe fecundity compensation prior to parasitic castration. Rearing the offspring produced at different times following the parent’s parasite exposure, our objective is to characterize the resistance of offspring by challenging them with the same pathogen when they reach the same age. Offspring resistance was measured as infection prevalence, while parasite virulence was estimated by cercarial output (the parasite’s reproductive success) (Nowak and May 1994). We hypothesize that there will be evidence of TGIP, which would be revealed by a lower infection prevalence and/or a lower infection intensity in offspring from an infected parent. Furthermore, we predict that there will be differential resistance in offspring depending on the time at which those offspring were produced in their parent’s infection. We expect offspring laid early in a parent’s infection (before fecundity compensation and subsequent parasite development) will demonstrate the highest resistance, evidenced by a low prevalence and virulence, due to a parent’s ability to allocate more resources to offspring before the host life history response and then the infection take their toll.

The NMRI strain of *S. mansoni* was maintained in our laboratory utilizing *B. glabrata* snails and Balb/c mice. All snails were grown under controlled laboratory conditions with a twelve-hour light and dark cycle and a room temperature of 25°C. Snails were housed in jars of well water and provided fresh lettuce *ab libitum* for the duration of the experiments.

A total of 198 snails between 9-15mm in length were initially divided into an experimental parent group of 150, and a control parent group of 48. Snails from the experimental group were individually exposed to five miracidia, and the control group snails were sham exposed in well plates, where they remained for twelve-hours before being transferred to individual jars of 225 mL volume. Styrofoam was placed in each jar as an egg-laying substrate. Egg masses were collected from the Styrofoam and separated out for ten weeks at discrete weekly intervals (Figure 1). Fecundity was recorded weekly, taken as the number of egg masses laid per individual each week (Gleichsner, Cleveland, and Minchella 2016). Egg collection coincided with weekly water changes, as parents were moved to new jars of well water. After five weeks, non-laying and dead parents were removed from the experiment, culling 19 parents from the experimental group and 8 parents from the control group. Parental infection was verified by the release of cercariae in the remaining experimental group 5-6 weeks following parasite exposure. Snails were placed in well-plates and exposed to a light for 1 hour to stimulate cercariae emergence (Lewis et al. 1986). The proportion of experimental parents infected (n=131) was recorded as a measure of prevalence. Uninfected snails in the experimental group were removed and the control group was randomly culled to 25 individuals. Offspring collected from removed parents were discarded from the experiment.

**Figure 1.**
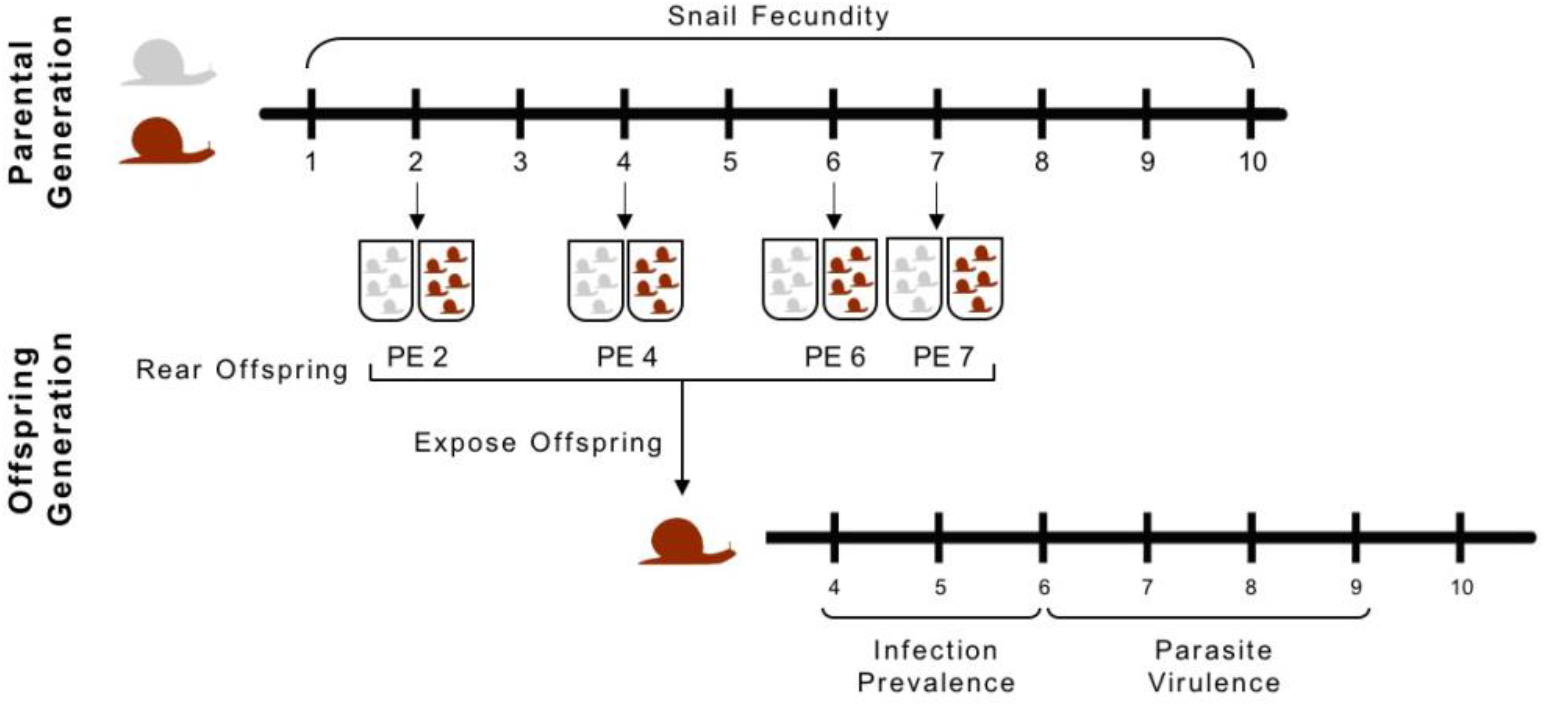
Experimental Design. Snail fecundity for infected (red) and uninfected control (gray) parental generation was recorded for 10 weeks post exposure (PE). Offspring from weeks 2, 4, 6, and 7 PE were collected and reared from both control and infected parents. The offspring generation was then exposed to *S. mansoni* at 12 weeks. Infection prevalence (at weeks 4-6) and parasite virulence (at weeks 6-9) were recorded for of the offspring generation.

Offspring were differentiated by both their parent ID and week of collection following the parent’s parasite exposure, which was designated post exposure (PE). Offspring from both control and infected parents were collected post exposure and were reared to be infected (Figure 1). Offspring from the same parent and week PE were housed together as a set for the duration of the experiment. Two weeks following their collection, hatched offspring were moved to new jars with 450 mL volume of well water. At ten weeks, 4-7 offspring from each set were randomly selected to continue and moved to a jar of fresh well water. At twelve weeks, 4-6 offspring from each set were individually exposed to 5 miracidia in well plates. Unexposed offspring were discarded. The exposed offspring continued to be housed together for the duration of the experiment with water changes occurring two times a week.

Infection was verified by cercariae shedding 4-6 weeks after parasite exposure. Prevalence was recorded as the proportion of infected offspring in each set 6 weeks following parasite exposure. Virulence was quantified by the cercariae output 6-10 weeks following parasite exposure (Figure 1). Virulence was only quantified in the groups of offspring with a minimum of 3 infected snails. If more than 3 offspring were infected, 3 infected individuals were randomly chosen, the unselected offspring were discarded. The selected snails were placed individually into well plates with 10 milliliters of well water, where they were exposed to light for 1 hour to induce cercariae emergence. 333 microliter aliquots were taken from each of the 3 wells to total a 999-microliter aliquot. Parasite virulence was calculated by averaging the total number of cercariae collected from the three aliquots.

Utilizing the offspring prevalence data from 6 weeks following exposure, a two-proportion z test was applied to test whether infection prevalence was significantly different in offspring based on parental infection status. Virulence data was Box-Cox transformed to satisfy the statistical assumptions for analysis. Our analysis showed that an exponent of 0.28 provided the best fit to a normal distribution. Data were back transformed for use in figures. Virulence data were analyzed using a general linear mixed model (GLMM) with parental treatment groups as a fixed factor and parental ID and offspring weeks post exposure as random effects. When analyzing virulence in the offspring associated with parental age, we used a GLMM with parents’ week post exposure as a fixed effect and parental ID, offspring week post exposure, and parental infection status as random effects.

A significant increase in reproduction was observed in the infected parent group compared to controls at three weeks following parasite exposure. A decrease in fecundity is visible in the infected parent group as time progresses and the infection becomes patent (Figure 2).

**Figure 2.**
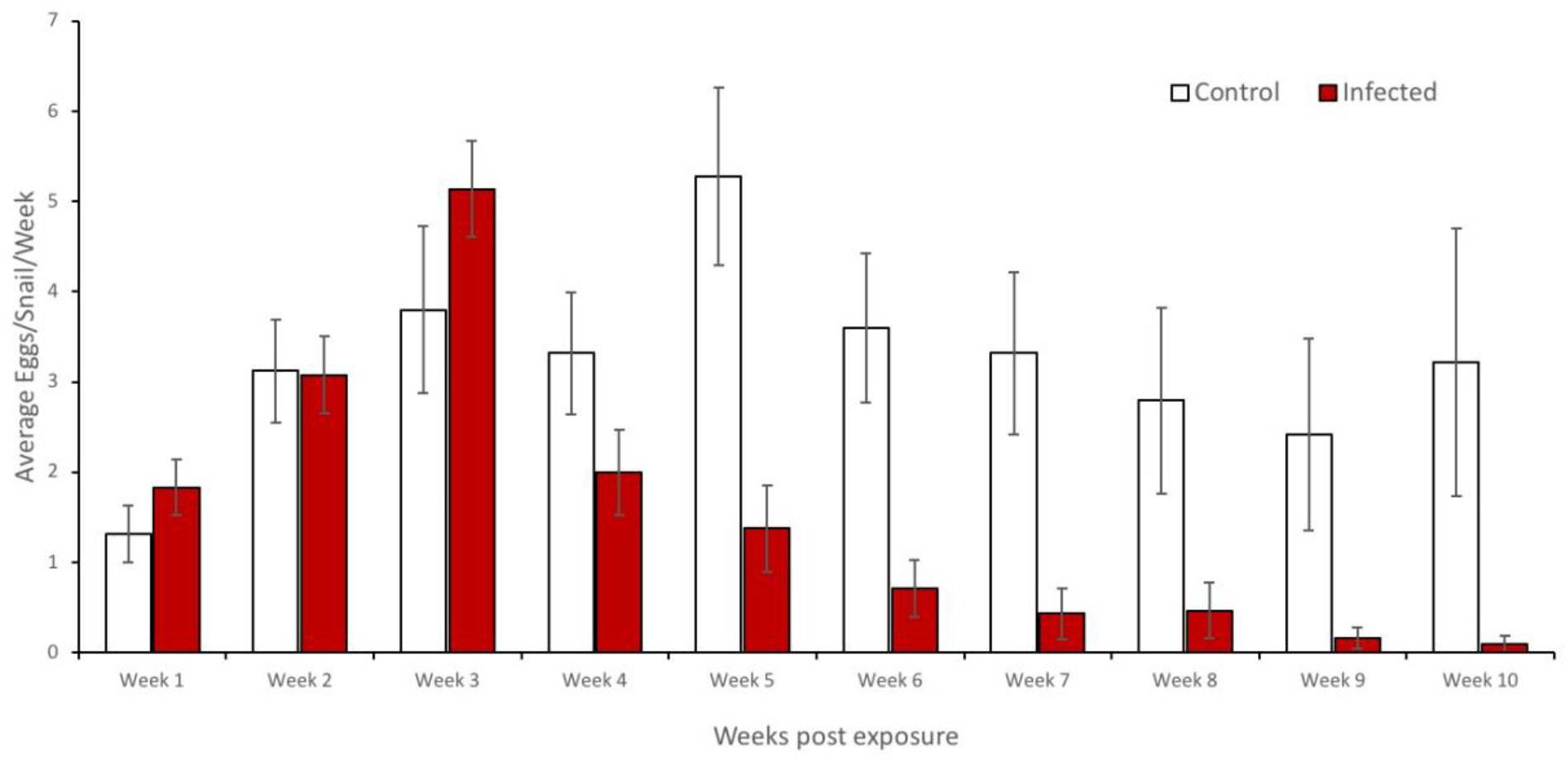
Snail fecundity of the parental generation over 10 weeks. Fecundity was calculated as the average reproductive output (number of egg masses) from each snail per week.

Offspring produced by infected parents two weeks post exposure have a significantly higher infection prevalence than offspring produced at the same timepoint from the control parents (χ2 = 16.906, p < 0.001). However, no other week PE exhibited a significant difference in infection prevalence between offspring of infected and control parents (Week 4 PE χ2 = 0.156, p = 0.693; Week 6 PE χ2 = 0.059, p = 0.809; Week 7 PE χ2 = 0.547, p = 0.460). Parasite virulence did not differ between infected offspring from infected parents or control parents for any time period post exposure (Week 4 PE F_1, 221_ = 1.315, p = 0.253; week 6 PE F_1, 124_ = 2.659, p = 0.106; week 7 PE F_1, 72_ = 2.344, p = 0.130; Figure 3)

**Figure 3:**
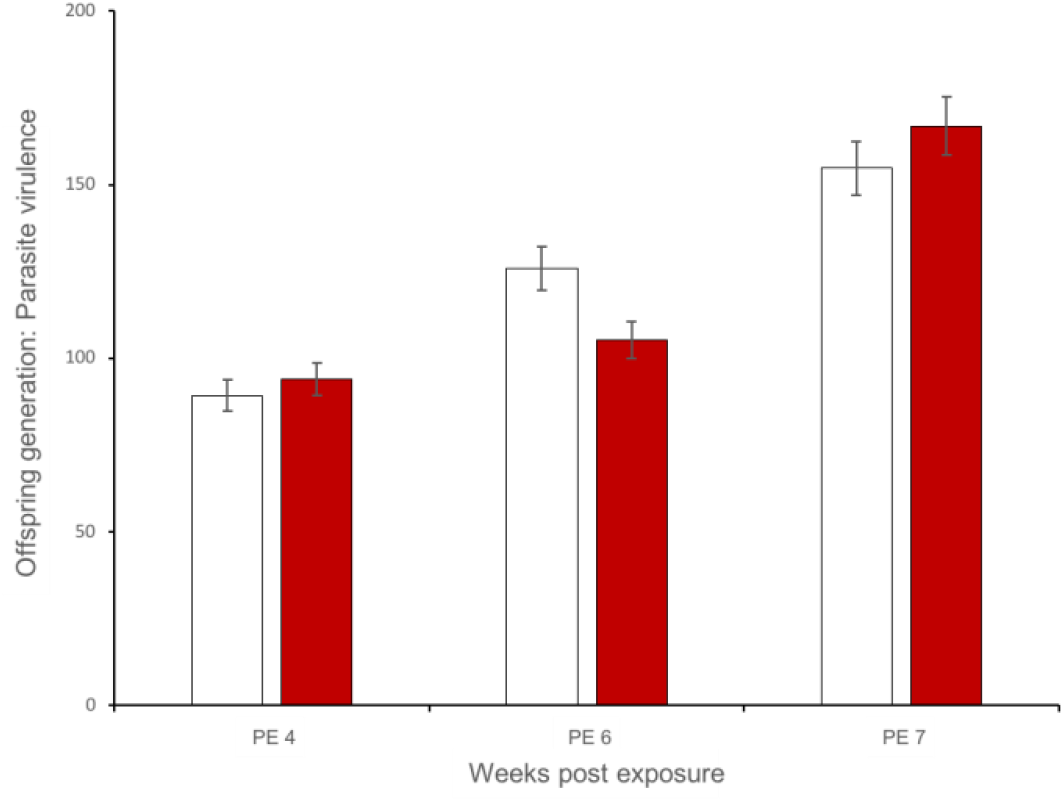
Parasite virulence (average number of cercariae) of offspring reared from parental generation at weeks 4, 6, and 7 post exposure with relation to parent infection status (control vs. infected). Virulence was not significantly different based on parental infection status for any week post exposure (PE 4 F_1, 221_ = 1.315, p = 0.253; PE 6 F_1, 124_ = 2.659, p = 0.106; PE 7 F_1, 72_ = 2.344, p = 0.130).

Parental investment theory predicts that if organisms can recognize an environmental challenge, they can make context-based investments into their offspring with non-genetic resources (Rosengaus et al. 2017). In *S. mansoni* infections of *B. glabrata,* a decrease in reproduction is obligate, as snails lose reproductive ability due to parasitic castration. There is also a period after parasite exposure and before castration that infected snails can selectively allocate resources to maximize their reproduction. Fecundity compensation represents a life history strategy to maximize reproductive output. In this study, fecundity compensation was observed in the third week following infection (Figure 1). However, accessing fecundity alone does not forecast the adaptiveness of offspring to parasitic challenges (Stahlschmidt et al. 2013).

Offspring adaptiveness in a parasite environment can be estimated by their resistance to parasitic infection. Evidence for an immune priming response in *B. glabrata* already challenges the traditional assumption that invertebrates lack immune memory (Portela et al. 2013) by indicating that resistance can be generated within generations. Furthermore, examples of transgenerational immune priming in an array of invertebrates (Mitchell and Read 2005; Baron et al. 2013; Rosengaus et al. 2017) are indicative that inherited invertebrate immunity can extend beyond genetic inheritance. TGIP can occur via deliverance of active immune components to offspring or by heightened endogenous immune function (Roth et al. 2018). TGIP in snails parasitized with *S. mansoni* could suggest that *S. mansoni* resistance is carried to offspring by processes beyond genetic changes. However, the results of this study do not provide support for TGIP in this system. In contrast, offspring from infected parents 2 weeks PE exhibited a significantly higher infection prevalence than offspring from uninfected parents. In fact, offspring from both infected and uninfected parents in week 2 demonstrated significantly lower prevalence than offspring at all other time periods. This result suggests that increased parasite resistance in offspring may be generated by healthier parents. Assessing infection virulence could provide a stronger reflection of whether PE 2 offspring demonstrated increased resistance. If the virulence of infections was lower in offspring from infected parents, this could indicate that the hosts had a more resistant/tolerant relationship with the parasite. Unfortunately, we were unable to assess virulence in week 2. The prevalence of infection was so low in the offspring from the control parent PE 2 group, that the number of individuals in the samples did not meet our standards to be analyzed for virulence. Virulence data were collected for offspring from weeks 4, 6, and 7 PE.

There was no indication of increased resistance from offspring of infected parents at weeks 4, 6, and 7 post exposure. Parental infection status did not confer any significant difference in virulence or infection prevalence within those offspring groups (Figure 3). Offspring from both the infected and control groups at the PE2 timepoint exhibited a significantly lower prevalence than offspring from all other timepoints. Differences in the prevalence of the PE 2 offspring may be a result of a healthier parent early in an infection. The results of this study do not support TGIP even though immune priming has been observed in this system (Portela et al. 2013). TGIP may not be an adaptive response in this host-parasite system. TGIP only benefits offspring if it is predictive of challenges those offspring will experience (Roth et al. 2018). The low prevalence of *S. mansoni* infections in natural snail populations, (often much less than five percent), may make the cost of implementing TGIP exceed the potential benefit (Minchella 1985).

We acknowledge the members of the Minchella lab group for their assistance in experimental set-up and data collection. We would also like to thank T.J. Vannatta who provided valuable support in the statistical analysis. *B. glabrata* snails provided by the NIAID Schistosomiasis Resource Center of the Biomedical Research Institute (Rockville, MD) through NIH-NIAID Contract HHSN272201700014I for distribution through BEI Resources. O.J. Lockyear was supported by a Cable-Silkman Undergraduate Fellowship. S.O Gutierrez was supported by the National Science Foundation Graduate Research Fellowships Program.

We would like to thank the members of the Minchella lab group for their assistance in data collection and T.J. Vannatta who provided valuable support in the statistical analysis. O.J. Lockyear was supported by a Cable-Silkman Undergraduate Fellowship. S.O Gutierrez was supported by the National Science Foundation Graduate Research Fellowships Program.

